# quincunx: an R package to query, download and wrangle PGS Catalog data

**DOI:** 10.1101/2021.02.19.431997

**Authors:** Ramiro Magno, Isabel Duarte, Ana -Teresa Maia

## Abstract

**Motivation:** The Polygenic Score (PGS) Catalog is a recently established open database of published polygenic scores that, to date, has collected, curated, and made available 721 polygenic scores from over 133 publications. The PGS Catalog REST API is the only method allowing programmatic access to this resource.

**Results:** Here, we describe *quincunx*, an R package that provides the first client interface to the PGS Catalog REST API. *quincunx* enables users to query and quickly retrieve, filter and integrate metadata associated with polygenic scores, as well as polygenic scoring files in tidy table format.

**Availability:** *quincunx* is freely available under an MIT License, and can be accessed from https://github.com/maialab/quincunx.

## Introduction

For two decades, GWAS identified individual variants associated with risk for complex diseases, raising the hopes of a polygenic approach for disease prevention. However, until recently, integration of these results was challenging delaying its prompt application to the clinical setting. In 2020 alone, over 1,400 publications on polygenic risk scores (PGS) appeared in PubMed, raising the need for a standardised distribution of studies’ key data, assuring their wide evaluation and accurate use.

The Polygenic Score (PGS) Catalog, created in 2019, is a publicly available, manually curated, open database of PGS and relevant metadata, that responds to this need [1]. Its current release [date 2021-02-03] includes manually curated data from 133 publications and 721 PGS associated with 194 traits. Currently, there are three alternative ways to access the data: (i) the web graphical user interface (GUI); (ii) by downloading database dumps; and (iii) the recently implemented PGS Catalog representational state transfer (REST) application programming interface (API), released in [date 2020-06-03], which provides direct programmatic access to the database, being this the preferred method for batch analyses.

We developed *quincunx*, the first R package [2] to programmatically access the PGS Catalog REST API. This package provides a simple user-friendly interface for querying the most updated Catalog data, retrieve and map it to in-memory relational databases of tidy data tables, allowing its prompt integration with tidyverse packages for subsequent data transformation, visualisation and modelling [3, 4].

## Results

### Retrieving data from the PGS Catalog REST API

The PGS Catalog REST API is an EBI service hosted at https://www.pgscatalog.org/rest/. The REST API uses hypermedia with resource responses following the OpenAPI Specification (https://swagger.io/docs/specification/about/). Response data is provided as hierarchical data in JSON format and can be paginated, i.e., split into multiple responses (https://www.pgscatalog.org/rest/).

To ease the conversion from the hierarchical to the relational tabular format — the preferred format for data analysis in R [4] — we developed a set of retrieval functions (Fig. 1A). Since the REST API data is organised around five core data entities — *Polygenic Scores*, *PGS Publications*, *PGS Sample Sets*, *PGS Performance Metrics* and *EFO traits*— we implemented five corresponding retrieval functions that encapsulate the technical aspects of resource querying and format conversion: `get_scores()`, `get_publications()`, `get_sample_sets()`, `get_performance_metrics` and `get_traits()` (Fig. 1A). These functions simplify the querying of PGS Catalog entities, by providing a complete and consistent interface to the Catalog. For example, to query for *scores*, the user needs only to know the function `get_scores()`, whereas the REST API itself exposes three separate resource URL endpoints for *scores* with different querying parameters. Moreover, the user can choose directly the arguments of the retrieval functions from any number of available search criteria exposed by the REST API (Fig. 1B). All arguments are vectorised, meaning that multiple queries are promptly available from a single function call. Results obtained from multiple queries can be combined with the logical operators OR or AND using the `set_operation` parameter. If `set_operation` is set to OR (default behaviour), results are collated while removing duplicates, if any. If `set_operation` is set to AND, only entities that concomitantly match all search criteria are returned. If finer control is needed on combining results, the following functions can be used: `bind()`, `union()`, `intersect()`, `setdiff()`, and `setequal()`. These are S4 methods that work with the S4 classes created in *quincunx*. An example of a case study (in tutorial style) can be found in Additional file 1.

**Figure 1.**
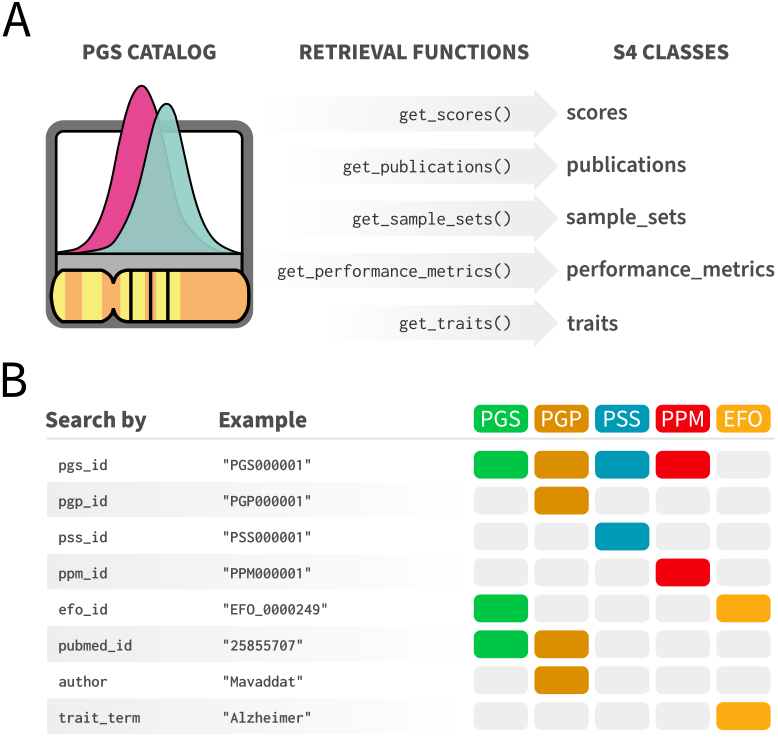
*quincunx* retrieval functions. **(A)** Functions for retrieving data from the PGS Catalog: `get_scores()`, `get_publications()`, `get_sample_sets()`, `get_performance_metrics()` and `get_traits()`. **(B)** *quincunx* search criteria (function parameters) to be used with retrieval functions. Coloured boxes indicate which entities can be retrieved by each search criteria.

### Representation of PGS Catalog entities

All S4 classes share the same design principles that make them relational databases: (i) each slot corresponds to a table (dataframe in R); (ii) the first slot corresponds to the main table that lists observations of the respective PGS Catalog entity, e.g., *scores*; and (iii) all tables have a primary key, the identifier of the respective PGS Catalog entity: `pgs_id`, `pgp_id`, `pss_id`, `ppm_id` or `efo_id`. For easy consultation of the variables present in the retrieved data tables, we provide a cheatsheet (Additional file 2); and for detailed descriptions, the user can issue the following help commands to open the respective help pages for each class: class?scores, class?publications, class?sample_sets, class?performance_metrics or class?traits.

### Improvements & Limitations

Compared to the exposed REST API, we have improved data accessibility in *quincunx* in several ways. Firstly, we harmonised the nomenclature of the variables in tidy tables with the nomenclature used by the GWAS Catalog [5], namely for the variables that are also used by the R package *gwasrapidd* [6]: an analogous R package that provides access to the GWAS Catalog REST API. This permits a frictionless wrangling of variables between the two R packages, allowing crosstalk between the data from the two Catalogs. Secondly, by recognising that in some cases the values of a variable are provided in its name and not in its value (a case of untidy data), we decided to perform the required refactoring to make those variables explicit columns in the relational tables, thus making the data more analysis friendly. For example, the *stage* of a sample comes implicitly coded in the JSON keys `samples_variants` and `samples_training` and are mapped in *quincunx* to the variable stage, with values ‘‘discovery’’ and ‘‘training’’, respectively. Additionally, the PGS Catalog REST API does not offer specific endpoints allowing direct mapping between the PGS entities, as this information is deeply nested in the hierarchical structure of the JSON responses. *quincunx* facilitates the retrieval of relationships between entities, by providing a set of mapping functions based on the entities’ identifiers, e.g., `pgs_to_pgp, pgp_to_ppm(), ppm_to_pss()`, including mapping (when applicable) from PGS scores to GWAS studies: `pgs_to_study() and study_to_pgs()` (see online documentation for the complete list). Finally, *quincunx* provides a set of helper functions to easily browse linked web resources, such as PubMed (`open_in_pubmed()`), dbSNP (`open_in_dbsnp()`), and the PGS Catalog Web interface itself (`open_in_pgs_catalog()`).

Despite the availability of some R software packages, e.g. *bigsnpr* [7], *RápidoPGS* [8], or *SummaryLasso* [9], that allow the application of polygenic scores to particular datasets, i.e. for analyses downstream of *quincunx* data retrieval, the most popular software tools for these calculations, e.g. PRSice [10, 11], *LDpred* [12], *PRS-CS* [13], *JAMPred* [14], *lassosum* [15], *PLINK* [16, 17], do not run in R. This could present an obstacle to pursuing a full PGS analysis within the same R framework (using the PGS scoring files), therefore delaying the process of polygenic score application (for an overview of the currently available methods, please see [18]).

## Conclusion

We have developed the first R client for the PGS Catalog REST API, thus greatly facilitating the programmatic access to the database. The main advantages of *quincunx* are: (i) providing a simple user-friendly interface to the REST API, allowing the programmatic querying of the most updated data from within R; (ii) the retrieval of the data in an analysis friendly format, with tidy data representations of the PGS entities, i.e., of *scores*, *publications*, *sample sets*, *performance metrics* and *traits* in the form of in-memory relational databases; (iii) allowing the automatic retrieval of polygenic scoring files from the PGS Catalog FTP server, making the data immediately available for analysis in R (as an extra feature not available via the REST API); and (iv) dedicated functions to export the retrieved objects to Excel (.xlsx) format for data inspection and sharing outside of R. *quincunx* is a package that will greatly improve the research community’s ability for data mining within R, therefore accelerating the evaluation and subsequent application of published and manually curated polygenic scores.

### Availability and requirements

- **Project name**: quincunx.
- **Project home page**: https://github.com/maialab/quincunx.
- **Operating system(s)**: Platform independent.
- **Programming language**: R.
- **Other requirements**: None.
- **License**: MIT.
- **Any restrictions to use by non-academics**: None.

## Supporting information

Additional File 1 - Example Study Case

Additional File 2 - quincunx cheatsheet

## Author’s contributions

RM & ID devised and wrote the package, and wrote the manuscript. ATM supervised the project and wrote the manuscript. All authors read and approved the final manuscript.

## Acknowledgements

We would like to thank the PGS Catalog team, particularly Samuel Lambert and Laurent Gil, for feedback and for supporting the development of *quincunx*. We also would like to thank the members of the Cancer Functional Genomics lab for user experience feedback, and UAIC-UAlg for administrative support.

## Additional Files

### Additional file 1 — Example of a case study

- **File name**: `additional_file_1.pdf`.
- **File format**: Portable Document Format (PDF).
- **Title**: Example Study Case.
- **Description**: Example of a study case exploring the PGS scores by Mavaddat et al. (2018).

### Additional file 2 — quincunx cheatsheet

- **File name**: `additional_file_2.pdf`
- **File format**: Portable Document Format (PDF)
- **Title**: quincunx cheatsheet
- **Description**: Additional file 2 contains an infographics: quincunx cheatsheet.

