## Additional File 1 - Example Study Case for "quincunx: an R package to query, download and wrangle PGS Catalog data"

### Exploring PGS scores by Mavaddat et al. (2018)

#### Introduction

Let us assume that you are a breast cancer researcher, and that you are interested in studying the screening and prevention of this disease. Now, imagine you have just recently noticed a new publication that claims to have developed a set of new polygenic risk scores that are both powerful and reliable predictors of breast cancer risk, e.g., Mavaddat et al. (2018) (https://doi.org/10.1016/j.ajhg.2018.11.002).

Perhaps now you would like to investigate a bit more about these new polygenic risk scores to assess their potential application. You know that the performance of these scores is dependent on various aspects, such as study design, participant demographics, case definitions, and covariates that have been adjusted for. In general, to access this information, you will have to carefully read the paper searching for these details, and usually get them from the supplementary material, with all the extra effort it takes. However, if these scores have already been indexed and manually curated by the PGS Catalog (https://www.pgscatalog.org/) team, then you can benefit by using this free and open resource to quickly gather relevant data about these scores. And if you happen to be an R user, then you can use the R Package *quincunx* and its retrieval functions to fetch the polygenic score metadata associated with the publication of interest from the PGS Catalog REST API server. This is what we will be doing next.

#### Finding polygenic risk scores by publication

##### Searching for publications by author name

To check if we are fortunate enough to have the polygenic risk scores for breast cancer (Mavaddat et al. (2018) (https://doi.org/10.1016/j.ajhg.2018.11.002)) in the PGS Catalog, we start by loading the quincunx package and by searching for this publication in the PGS Catalog. To do this we will use the function `get_publications()` and start by searching by the first author's last name, i.e. "Mavaddat":

```
library(quincunx)
library(dplyr)

pub_by_mavaddat <- get_publications(author = 'Mavaddat') # Not case sensitive,
                                                          # so "mavaddat" would
                                                          # have worked just fine.
```

When you ask the PGS Catalog for publications, it returns an S4 `publications` object with two tables (slots): `publications` and `pgs_ids`. You can access these tables with the `@` operator.

The `publications` table (data frame) contains all the publications added to the PGS Catalog that contain an author whose last name is "Mavaddat":

```
pub_by_mavaddat@publications
#> # A tibble: 6 x 8
#>   pgp_id pubmed_id publication_date publication title author_fullname doi
#>   <chr>  <chr>     <date>           <chr>       <chr> <chr>           <chr>
#> 1 PGP00… 25855707  2015-04-08       J Natl Can… Pred… Mavaddat N      10.1…
#> 2 PGP00… 30554720  2018-12-13       Am J Hum G… Poly… Mavaddat N      10.1…
#> 3 PGP00… 28376175  2017-07-01       J Natl Can… Eval… Kuchenbaecker … 10.1…
#> 4 PGP00… 33022221  2020-10-05       Am J Hum G… Brea… Kramer I        10.1…
#> 5 PGP00… 32737321  2020-07-31       Nat Commun  Euro… Ho WK           10.1…
#> 6 PGP00… 32665703  2020-07-15       Genet Med   Poly… Barnes DR       10.1…
#> # … with 1 more variable: authors <chr>
```

Seemingly there are 6 publications indexed in the PGS Catalog. Each publication added to the PGS Catalog gets an unique identifier. This identifier is in the first column ( `pgp_id` ) of the `publications` table. The number of publications can be obtained by using the function `n()` on the S4 object `pub_by_mavaddat`, or by asking directly the number of rows of the `publications` table:

```
quincunx::n(pub_by_mavaddat) # Here we use quincunx::n() instead of n() to avoid name
space collisions with dplyr::n().
#> [1] 6
nrow(pub_by_mavaddat@publications)
#> [1] 6
```

Moreover, we can see that Nasim Mavaddat is not the first author in all these publications because her name is not always present in the column `author_fullname` (which contains the name of the first author only). If you want to know all the authors you can access the column `authors` (contains a string of comma separated author names).

Now, to identify our publication of interest amongst these 6 publications, we can check, for example, the PubMed identifier of Mavaddat et al. (2018) (https://doi.org/10.1016/j.ajhg.2018.11.002) (column `pubmed_id`), or check other information such as journal name (column `publication`), the publication title (column `title`) or the publication date (column `publication_date`). The most unambiguous approach is to use the PubMed identifier. The PubMed identifier for Mavaddat et al. (2018) (https://doi.org/10.1016/j.ajhg.2018.11.002) is `"30554720"`. So the publication of interest corresponds to the one returned in row number 2 of the `publications` table:

```
publication_row_index <- which(pub_by_mavaddat@publications$pubmed_id == "30554720")
publication_row_index
#> [1] 2
```

To focus now on this publication and filter out the others, we can subset the `publications` object `pub_by_mavaddat`, e.g., by position, and get a new object with both tables (`publications` and `pgs_ids`) filtered for only this publication:

```
mavaddat2018 <- pub_by_mavaddat[publication_row_index]
mavaddat2018
#> An object of class "publications"
#> Slot "publications":
#> # A tibble: 1 x 8
#>   pgp_id pubmed_id publication_date publication title author_fullname doi
#>   <chr>  <chr>     <date>           <chr>       <chr> <chr>           <chr>
#> 1 PGP00… 30554720  2018-12-13       Am J Hum G… Poly… Mavaddat N      10.1…
#> # … with 1 more variable: authors <chr>
#>
#> Slot "pgs_ids":
#> # A tibble: 15 x 3
#>    pgp_id    pgs_id    stage
#>    <chr>     <chr>     <chr>
#>  1 PGP000002 PGS000004 development
#>  2 PGP000002 PGS000005 development
#>  3 PGP000002 PGS000006 development
#>  4 PGP000002 PGS000007 development
#>  5 PGP000002 PGS000008 development
#>  6 PGP000002 PGS000009 development
#>  7 PGP000002 PGS000001 evaluation
#>  8 PGP000002 PGS000002 evaluation
#>  9 PGP000002 PGS000003 evaluation
#> 10 PGP000002 PGS000004 evaluation
#> 11 PGP000002 PGS000005 evaluation
#> 12 PGP000002 PGS000006 evaluation
#> 13 PGP000002 PGS000007 evaluation
#> 14 PGP000002 PGS000008 evaluation
#> 15 PGP000002 PGS000009 evaluation
```

We see now that the `publications` table contains one entry only, and that the PGS Catalog identifier assigned to this publication is PGP000002:

```
mavaddat2018@publications$pgp_id
#> [1] "PGP000002"
```

In the `pgs_ids` we see that there are several PGS scores associated with publication PGP000002. Besides listing the score identifiers ( `pgs_id` ), it also includes the `stage` column that annotates the polygenic score relative to the PGS construction process (check page 2 of the quincunx cheatsheet (https://github.com/ramiromagno/cheatsheets/blob/master/quincunx/quincunx_cheatsheet.pdf) for a visual illustration).

#### Searching for publications by PubMed ID

An alternatively, albeit more direct, route to get this publication by Mavaddat et al. (2018) (https://doi.org/10.1016/j.ajhg.2018.11.002) could have been to query for publications directly by the corresponding PubMed ID (30554720):

```
pub_by_pmid_30554720 <- get_publications(pubmed_id = '30554720')
pub_by_pmid_30554720
#> An object of class "publications"
#> Slot "publications":
#> # A tibble: 1 x 8
#>   pgp_id pubmed_id publication_date publication title author_fullname doi
#>   <chr>  <chr>     <date>           <chr>       <chr> <chr>           <chr>
#> 1 PGP00… 30554720  2018-12-13       Am J Hum G… Poly… Mavaddat N      10.1…
#> # … with 1 more variable: authors <chr>
#>
#> Slot "pgs_ids":
#> # A tibble: 15 x 3
#>    pgp_id    pgs_id    stage
#>    <chr>     <chr>     <chr>
#>  1 PGP000002 PGS000004 development
#>  2 PGP000002 PGS000005 development
#>  3 PGP000002 PGS000006 development
#>  4 PGP000002 PGS000007 development
#>  5 PGP000002 PGS000008 development
#>  6 PGP000002 PGS000009 development
#>  7 PGP000002 PGS000001 evaluation
#>  8 PGP000002 PGS000002 evaluation
#>  9 PGP000002 PGS000003 evaluation
#> 10 PGP000002 PGS000004 evaluation
#> 11 PGP000002 PGS000005 evaluation
#> 12 PGP000002 PGS000006 evaluation
#> 13 PGP000002 PGS000007 evaluation
#> 14 PGP000002 PGS000008 evaluation
#> 15 PGP000002 PGS000009 evaluation
```

To get an overview of the possible search criteria for `get_publications` you can use the help function within R.

```
?get_publications

# or alternatively
help("get_publications")
```

### From publication to polygenic risk scores

Now that we have found that our publication of interest exists in the PGS Catalog, with identifier PGP000002, we can check now which polygenic risk scores are annotated in the Catalog. Polygenic scores (PGS) in the PGS Catalog are indexed by an unique accession identifier of the form: “PGS000000” (“PGS” followed by six digits).

To get all PGS identifiers associated with Mavaddat’s publication we turn to the second table `pgs_ids` that maps publication identifiers (PGP) to PGS identifiers:

```
pub_by_pmid_30554720@pgs_ids
#> # A tibble: 15 x 3
#>    pgp_id    pgs_id    stage
#>    <chr>     <chr>     <chr>
#>  1 PGP000002 PGS000004 development
#>  2 PGP000002 PGS000005 development
#>  3 PGP000002 PGS000006 development
#>  4 PGP000002 PGS000007 development
#>  5 PGP000002 PGS000008 development
#>  6 PGP000002 PGS000009 development
#>  7 PGP000002 PGS000001 evaluation
#>  8 PGP000002 PGS000002 evaluation
#>  9 PGP000002 PGS000003 evaluation
#> 10 PGP000002 PGS000004 evaluation
#> 11 PGP000002 PGS000005 evaluation
#> 12 PGP000002 PGS000006 evaluation
#> 13 PGP000002 PGS000007 evaluation
#> 14 PGP000002 PGS000008 evaluation
#> 15 PGP000002 PGS000009 evaluation
```

Please note that there are 9 unique scores, both from the *development* and the *evaluation* stages, meaning that this paper published new polygenic scores (*development* stage), and tested them (*evaluation* stage). But this paper has also evaluated 3 other polygenic scores previously developed (and firstly published in another publication by the same author).

**This distinction between stages is important because when we query the database for the scores from all the pgp_ids present in this publication, only the newly developed ones (from the *development stage*) will be retrieved.** (See below: section about the `get_scores()` function).

```
# Newly published PGS scores (development stage)
pub_by_pmid_30554720@pgs_ids %>% dplyr::filter(stage == "development")
#> # A tibble: 6 x 3
#>   pgp_id    pgs_id    stage
#>   <chr>     <chr>     <chr>
#> 1 PGP000002 PGS000004 development
#> 2 PGP000002 PGS000005 development
#> 3 PGP000002 PGS000006 development
#> 4 PGP000002 PGS000007 development
#> 5 PGP000002 PGS000008 development
#> 6 PGP000002 PGS000009 development

# All PGS scores evaluated (evaluation stage)
pub_by_pmid_30554720@pgs_ids %>% dplyr::filter(stage == "evaluation")
#> # A tibble: 9 x 3
#>   pgp_id    pgs_id    stage
#>   <chr>     <chr>     <chr>
#> 1 PGP000002 PGS000001 evaluation
#> 2 PGP000002 PGS000002 evaluation
#> 3 PGP000002 PGS000003 evaluation
#> 4 PGP000002 PGS000004 evaluation
#> 5 PGP000002 PGS000005 evaluation
#> 6 PGP000002 PGS000006 evaluation
#> 7 PGP000002 PGS000007 evaluation
#> 8 PGP000002 PGS000008 evaluation
#> 9 PGP000002 PGS000009 evaluation
```

If we knew, before hand, that PGP000002 was associated with Mavaddat's publication, we could have also taken advantage of the neat function `pgp_to_pgs()` to quickly get all the associated polygenic score ids:

```
pgp_to_pgs('PGP000002')
#> # A tibble: 15 x 3
#>    pgp_id    pgs_id    stage
#>    <chr>     <chr>     <chr>
#>  1 PGP000002 PGS000004 development
#>  2 PGP000002 PGS000005 development
#>  3 PGP000002 PGS000006 development
#>  4 PGP000002 PGS000007 development
#>  5 PGP000002 PGS000008 development
#>  6 PGP000002 PGS000009 development
#>  7 PGP000002 PGS000001 evaluation
#>  8 PGP000002 PGS000002 evaluation
#>  9 PGP000002 PGS000003 evaluation
#> 10 PGP000002 PGS000004 evaluation
#> 11 PGP000002 PGS000005 evaluation
#> 12 PGP000002 PGS000006 evaluation
#> 13 PGP000002 PGS000007 evaluation
#> 14 PGP000002 PGS000008 evaluation
#> 15 PGP000002 PGS000009 evaluation
```

### Polygenic Scores published by Mavaddat *et al.* (2018)

To dive into the metadata about these polygenic scores, we use the quincunx function `get_scores()`:

```
mavaddat2018_scores <- get_scores(pubmed_id = "30554720")
slotNames(mavaddat2018_scores)
#> [1] "scores"       "publications" "samples"      "demographics" "cohorts"
#> [6] "traits"
```

This returns the S4 object `scores` which contains six tables (slots): scores, publications, samples, demographics, cohorts, traits.
You can quickly check all variables from each table by consulting quincunx cheatsheet (https://github.com/ramiromagno/cheatsheets/blob/master/quincunx/quincunx_cheatsheet.pdf).
A description of each variable is annotated in the `scores` object help page that can be accessed with `class?scores`.

### Polygenic Scores Metadata

#### scores object | scores table

The S4 scores object `mavaddat2018_scores` starts with a table named `scores` that lists each score in one row. All scores are identified by a PGS identifier ( `pgs_id` column). Note that, as previously explained, only the polygenic scores newly developed in this publication (*developmental stage*) are retrieved, and not all the PGS scores that were evaluated in this publication.

In addition, scores can have a name ( `pgs_name` column). This may be the name assigned by authors of the source publication, or a name assigned by a PGS Catalog curator in order to identify that particular score throughout the curating process.

```
mavaddat2018_scores@scores[c('pgs_id', 'pgs_name')]
#> # A tibble: 6 x 2
#>   pgs_id    pgs_name
#>   <chr>     <chr>
#> 1 PGS000004 PRS313_BC
#> 2 PGS000005 PRS313_ERpos
#> 3 PGS000006 PRS313_ERneg
#> 4 PGS000007 PRS3820_BC
#> 5 PGS000008 PRS3820_ERpos
#> 6 PGS000009 PRS3820_ERneg
```

From the PGS names we can already see the presence of the suffixes “ERpos” and “ERneg”, suggestive of specialized polygenic risk scores for estrogen-receptor positive and negative samples.

```
mavaddat2018_scores@scores['scoring_file']
#> # A tibble: 6 x 1
#>   scoring_file
#>   <chr>
#> 1 http://ftp.ebi.ac.uk/pub/databases/spot/pgs/scores/PGS000004/ScoringFiles/PGS…
#> 2 http://ftp.ebi.ac.uk/pub/databases/spot/pgs/scores/PGS000005/ScoringFiles/PGS…
#> 3 http://ftp.ebi.ac.uk/pub/databases/spot/pgs/scores/PGS000006/ScoringFiles/PGS…
#> 4 http://ftp.ebi.ac.uk/pub/databases/spot/pgs/scores/PGS000007/ScoringFiles/PGS…
#> 5 http://ftp.ebi.ac.uk/pub/databases/spot/pgs/scores/PGS000008/ScoringFiles/PGS…
#> 6 http://ftp.ebi.ac.uk/pub/databases/spot/pgs/scores/PGS000009/ScoringFiles/PGS…
```

The column `scoring_file` contains the URL for the FTP location containing the corresponding PGS scoring files. PGS scoring files are the text files provided by the PGS Catalog team containing the source data that you can use to compute polygenic scores for particular individuals, i.e. that allow you to apply these scores to your individual samples. Learn more about scoring files in `vignette("pgs-scoring-file")`. For a quick consultation of the file format of PGS scoring files you may also check the second page of quincunx cheatsheet (https://github.com/ramiromagno/cheatsheets/blob/master/quincunx/quincunx_cheatsheet.pdf).

As an additional feature, *quincunx* allows you to download the relevant PGS scoring files directly into R using the function `read_scoring_file()`, making the data immediately available in R for further analysis.

```
mavaddat2018_scores@scores[c('pgs_id', 'matches_publication')]
#> # A tibble: 6 x 2
#>   pgs_id    matches_publication
#>   <chr>     <lgl>
#> 1 PGS000004 TRUE
#> 2 PGS000005 TRUE
#> 3 PGS000006 TRUE
#> 4 PGS000007 TRUE
#> 5 PGS000008 TRUE
#> 6 PGS000009 TRUE
```

The column `matches_publication` is a logical value indicating whether the published polygenic score is exactly the same as the one present in the PGS scoring file provided by the PGS Catalog. In this case all of the 6 scores are provided exactly as published (all values are "TRUE").

Other columns in the `scores` table hold relevant information.

```
mavaddat2018_scores@scores[c('pgs_name', 'pgs_method_name', 'pgs_method_params', 'n_variants', 'assembly')]
#> # A tibble: 6 x 5
#>   pgs_name    pgs_method_name              pgs_method_para… n_variants assembly
#>   <chr>       <chr>                        <chr>                 <int> <chr>
#> 1 PRS313_BC   Hard-Thresholding Stepwise F… p < 10^-5              313 GRCh37
#> 2 PRS313_ERp… Hard-Thresholding Stepwise F… p < 10^-5              313 GRCh37
#> 3 PRS313_ERn… Hard-Thresholding Stepwise F… p < 10^-5              313 GRCh37
#> 4 PRS3820_BC  LASSO penalized regression    p < 0.001             3820 GRCh37
#> 5 PRS3820_ER… LASSO penalized regression    p < 0.001             3820 GRCh37
#> 6 PRS3820_ER… LASSO penalized regression    p < 0.001             3820 GRCh37
```

For example, columns such as `pgs_method_name` and `pgs_method_params` provide extra details about the PGS development method. Finally, `n_variants` informs about the number of variants comprising each polygenic risk score, and `assembly` indicates the genome assembly version used.

#### scores object | publications table

The `scores` S4 object contains a table dedicated to the source publications used to collect the score(s) retrieved. In this case, it is not surprising that all PGS scores map to the same publication identifier, i.e., PGP000002, as that was our starting point.

```
mavaddat2018_scores@publications[c('pgs_id', 'pgp_id', 'publication_date', 'author_fu
llname')]
#> # A tibble: 6 x 4
#>   pgs_id    pgp_id    publication_date author_fullname
#>   <chr>     <chr>     <date>           <chr>
#> 1 PGS000004 PGP000002 2018-12-13       Mavaddat N
#> 2 PGS000005 PGP000002 2018-12-13       Mavaddat N
#> 3 PGS000006 PGP000002 2018-12-13       Mavaddat N
#> 4 PGS000007 PGP000002 2018-12-13       Mavaddat N
#> 5 PGS000008 PGP000002 2018-12-13       Mavaddat N
#> 6 PGS000009 PGP000002 2018-12-13       Mavaddat N
```

#### scores object | samples table

The third table (slot) of the `scores` S4 object pertains to the samples used for the development of the PGS scores.

There are a total of 15 columns with metadata details about each sample. Each row corresponds to one sample associated with the polygenic scores, and the combination of values of the first two columns, `pgs_id` and `sample_id`, uniquely identifies each sample in this table. All samples shown in the `samples` table of a `scores` object are annotated with a `stage`, that can take two values: `"discovery"` or `"training"`.
The `"discovery"` samples are typically used in determining the variants that are afterwards used for polygenic score development. These variants originate typically from Genome-Wide Association Studies (GWAS). Hence, these samples might be linked to the GWAS Catalog. If that is the case, this information is provided in the column `study_id`, indicating the GWAS Catalog accession identifier. You may find more information about these GWAS studies by using the function `gwasrapidd::get_studies()` from the gwasrapidd (https://rmagno.eu/gwasrapidd/) R package that we developed previously (described here (https://doi.org/10.1093/bioinformatics/btz605)).
The `"training"` samples are those that have been used for the training of a particular polygenic score. Together, these two stages (*discovery* and *training*) are referred to as *development*, in contrast to the later testing phase of the polygenic scores, i.e., the *evaluation* phase (or stage).
If this sounds confusing check our cheatsheet (https://github.com/ramiromagno/cheatsheets/blob/master/quincunx/quincunx_cheatsheet.pdf), section *PGS Construction Process*, second page.

```
dplyr::glimpse(mavaddat2018_scores@samples)
#> Rows: 18
#> Columns: 15
#> $ pgs_id                          <chr> "PGS000004", "PGS000004", "PGS000004"…
#> $ sample_id                       <int> 1, 2, 3, 1, 2, 3, 1, 2, 3, 1, 2, 3, 1…
#> $ stage                           <chr> "discovery", "training", "training", …
#> $ sample_size                     <int> 139274, 158648, 10444, 139274, 87368,…
#> $ sample_cases                    <int> NA, 88916, 5159, NA, 55391, 4233, NA,…
#> $ sample_controls                 <int> NA, 69732, 5285, NA, 31977, 926, NA, …
#> $ sample_percent_male             <dbl> NA, 0, 0, NA, 0, 0, NA, 0, 0, NA, 0, …
#> $ phenotype_description           <chr> NA, "Invasive breast cancer-affected"…
#> $ ancestry                        <chr> "European", "European", "European", "…
#> $ ancestry_description            <chr> "NR", NA, NA, "NR", NA, NA, "NR", NA,…
#> $ ancestry_country                <chr> "Canada, U.S., Australia, Belgium, Fr…
#> $ ancestry_additional_description <chr> NA, NA, NA, NA, NA, NA, NA, NA, NA, N…
#> $ study_id                        <chr> "GCST004988", NA, NA, "GCST004988", N…
#> $ pubmed_id                       <chr> "29059683", NA, NA, "29059683", NA, N…
#> $ cohorts_additional_description  <chr> NA, "Training Cohort", "Validation Co…
```

The PGS Catalog provides brief records of samples sizes (total number, number of cases, and number of controls):

```
mavaddat2018_scores@samples[1:6]
#> # A tibble: 18 x 6
#>    pgs_id    sample_id stage     sample_size sample_cases sample_controls
#>    <chr>         <int> <chr>           <int>        <int>           <int>
#>  1 PGS000004         1 discovery      139274           NA              NA
#>  2 PGS000004         2 training       158648        88916           69732
#>  3 PGS000004         3 training        10444         5159            5285
#>  4 PGS000005         1 discovery      139274           NA              NA
#>  5 PGS000005         2 training        87368        55391           31977
#>  6 PGS000005         3 training         5159         4233             926
#>  7 PGS000006         1 discovery      139274           NA              NA
#>  8 PGS000006         2 training        87368        15404           71964
#>  9 PGS000006         3 training         5159          926            4233
#> 10 PGS000007         1 discovery      139274           NA              NA
#> 11 PGS000007         2 training       158648        88916           69732
#> 12 PGS000007         3 training        10444         5159            5285
#> 13 PGS000008         1 discovery      139274           NA              NA
#> 14 PGS000008         2 training        87368        55391           31977
#> 15 PGS000008         3 training         5159         4233             926
#> 16 PGS000009         1 discovery      139274           NA              NA
#> 17 PGS000009         2 training        87368        15404           71964
#> 18 PGS000009         3 training         5159          926            4233
```

Perhaps, not so surprisingly, the percentage of male participants in the training samples is zero:

```
mavaddat2018_scores@samples[c('pgs_id', 'sample_id', 'stage', 'sample_percent_male')]
#> # A tibble: 18 x 4
#>    pgs_id    sample_id stage     sample_percent_male
#>    <chr>         <int> <chr>                   <dbl>
#>  1 PGS000004         1 discovery                  NA
#>  2 PGS000004         2 training                    0
#>  3 PGS000004         3 training                    0
#>  4 PGS000005         1 discovery                  NA
#>  5 PGS000005         2 training                    0
#>  6 PGS000005         3 training                    0
#>  7 PGS000006         1 discovery                  NA
#>  8 PGS000006         2 training                    0
#>  9 PGS000006         3 training                    0
#> 10 PGS000007         1 discovery                  NA
#> 11 PGS000007         2 training                    0
#> 12 PGS000007         3 training                    0
#> 13 PGS000008         1 discovery                  NA
#> 14 PGS000008         2 training                    0
#> 15 PGS000008         3 training                    0
#> 16 PGS000009         1 discovery                  NA
#> 17 PGS000009         2 training                    0
#> 18 PGS000009         3 training                    0
```

Also, information about the trait or disease studied and ancestry information can be accessed:

```
mavaddat2018_scores@samples[c('pgs_id', 'sample_id', 'stage', 'phenotype_description'
, 'ancestry')]
#> # A tibble: 18 x 5
#>    pgs_id    sample_id stage     phenotype_description           ancestry
#>    <chr>         <int> <chr>     <chr>                           <chr>
#>  1 PGS000004         1 discovery <NA>                            European
#>  2 PGS000004         2 training  Invasive breast cancer-affected European
#>  3 PGS000004         3 training  Invasive breast cancer-affected European
#>  4 PGS000005         1 discovery <NA>                            European
#>  5 PGS000005         2 training  ER-positive breast cancer cases European
#>  6 PGS000005         3 training  ER-positive breast cancer cases European
#>  7 PGS000006         1 discovery <NA>                            European
#>  8 PGS000006         2 training  ER-negative breast cancer cases European
#>  9 PGS000006         3 training  ER-negative breast cancer cases European
#> 10 PGS000007         1 discovery <NA>                            European
#> 11 PGS000007         2 training  Invasive breast cancer-affected European
#> 12 PGS000007         3 training  Invasive breast cancer-affected European
#> 13 PGS000008         1 discovery <NA>                            European
#> 14 PGS000008         2 training  ER-positive breast cancer cases European
#> 15 PGS000008         3 training  ER-positive breast cancer cases European
#> 16 PGS000009         1 discovery <NA>                            European
#> 17 PGS000009         2 training  ER-negative breast cancer cases European
#> 18 PGS000009         3 training  ER-negative breast cancer cases European
```

Again, not so surprisingly, all samples are of European ancestry, a bias recognized by the research community. You can find more details about the ancestry variable in the `ancestry_categories` column. These categories have been defined within the NHGRI-EBI GWAS Catalog framework. We provide these ancestry nomenclature in *quincunx* as a separate dataset named `ancestry_categories`. See `?ancestry_categories` more details. To quickly lookup the definition of the *European* ancestry:

```
# Quick look at the ancestries definitions
ancestry_categories
#> # A tibble: 17 x 3
#>    ancestry                    definition                       examples
#>    <chr>                       <chr>                            <chr>
#>  1 Aboriginal Australian       "Includes individuals who eithe… Martu Australia…
#>  2 African American or Afro-C… "Includes individuals who eithe… African America…
#>  3 African unspecified         "Includes individuals that eith… African, non-Hi…
#>  4 Asian unspecified           "Includes individuals that eith… Asian, Asian Am…
#>  5 Central Asian               "Includes individuals who eithe… Silk Road (foun…
#>  6 East Asian                  "Includes individuals who eithe… Chinese, Japane…
#>  7 European                    "Includes individuals who eithe… Spanish, Swedish
#>  8 Greater Middle Eastern (Mi… "Includes individuals who self-… Tunisian, Arab,…
#>  9 Hispanic or Latin American  "Includes individuals who eithe… Brazilian, Mexi…
#> 10 Native American             "Includes indigenous individual… Pima Indian, Pl…
#> 11 Not reported                "Includes individuals for which… <NA>
#> 12 Oceanian                    "Includes individuals that eith… Solomon Islande…
#> 13 Other                       "Includes individuals where an … Surinamese, Rus…
#> 14 Other admixed ancestry      "Includes individuals who eithe… <NA>
#> 15 South Asian                 "Includes individuals who eithe… Bangladeshi, Sr…
#> 16 South East Asian            "Includes individuals who eithe… Thai, Malay
#> 17 Sub-Saharan African         "Includes individuals who eithe… Yoruban, Gambian
```

#### scores object | demographics table

The demographics table usually lists demographic information about each sample. For this study this table is however empty, meaning that this information was either not available from Mavaddat's publication, or not included in the PGS Catalog.

```
mavaddat2018_scores@demographics
#> # A tibble: 0 x 11
#> # … with 11 variables: pgs_id <chr>, sample_id <int>, variable <chr>,
#> #   estimate_type <chr>, estimate <dbl>, unit <chr>, variability_type <chr>,
#> #   variability <dbl>, interval_type <chr>, interval_lower <dbl>,
#> #   interval_upper <dbl>
```

Nevertheless, the demographics variables, when present, are follow-up time, and age of study participants.

If you want to confirm that *quincunx* is retrieving exactly the same info as provided by the PGS Catalog web interface, you can always check this by showing online the metadata for your PGP publication of interest using the function `open_in_pgs_catalog`.

```
open_in_pgs_catalog('PGP000002', pgs_catalog_entity = 'pgp')
```

#### scores object | cohorts table

In the cohorts table you can check which cohorts are associated with which samples. Note that the unique identification of a sample is given by the combination of the values of the first two columns: `pgs_id`, and `sample_id`.
To learn more about the meaning of cohorts for the PGS Catalog, visit our check our cheatsheet (https://github.com/ramiromagno/cheatsheets/blob/master/quincunx/quincunx_cheatsheet.pdf), section *Cohorts, Samples and Sample Sets*, second page.

```
mavaddat2018_scores@cohorts
#> # A tibble: 558 x 4
#>    pgs_id    sample_id cohort_symbol cohort_name
#>    <chr>         <int> <chr>         <chr>
#>  1 PGS000004         2 ABCFS         Australian Breast Cancer Family Study
#>  2 PGS000004         2 MCCS          Melbourne Collaborative Cohort Study
#>  3 PGS000004         2 HMBCS         Hannover-Minsk Breast Cancer Study
#>  4 PGS000004         2 LMBC          Leuven Multidisciplinary Breast Centre
#>  5 PGS000004         2 MTLGEBCS      Montreal Gene-Environment Breast Cancer St…
#>  6 PGS000004         2 CGPS          Copenhagen General Population Study
#>  7 PGS000004         2 KBCP          Kuopio Breast Cancer Project
#>  8 PGS000004         2 OBCS          Oulu Breast Cancer Study
#>  9 PGS000004         2 CECILE        CECILE Breast Cancer Study
#> 10 PGS000004         2 BBCC          Bavarian Breast Cancer Cases and Controls
#> # … with 548 more rows
```

#### scores object | traits table

Finally, in the traits table, you have access to the traits (phenotypes) associated with these polygenic scores. In this study, all scores indicate “breast cancer” or one of its subtypes (column `trait`):

```
mavaddat2018_scores@traits[c('pgs_id', 'efo_id', 'trait')]
#> # A tibble: 6 x 3
#>   pgs_id    efo_id      trait
#>   <chr>     <chr>       <chr>
#> 1 PGS000004 EFO_0000305 breast carcinoma
#> 2 PGS000005 EFO_1000649 estrogen-receptor positive breast cancer
#> 3 PGS000006 EFO_1000650 estrogen-receptor negative breast cancer
#> 4 PGS000007 EFO_0000305 breast carcinoma
#> 5 PGS000008 EFO_1000649 estrogen-receptor positive breast cancer
#> 6 PGS000009 EFO_1000650 estrogen-receptor negative breast cancer
```

Compared to the author-reported trait (column `reported_trait` from table `scores`), the trait description in this table follows the controlled vocabulary of an ontology, i.e., the Experimental Factor Ontology (EFO) (https://www.ebi.ac.uk/efo/). This way, traits are described objectively. This is very useful for comparing trait data among different studies where different reported trait descriptions might have been used. For example, if you want now to know what other polygenic scores may be deposited in the PGS Catalog that also study breast cancer — namely, breast carcinoma, estrogen-receptor positive breast cancer, or estrogen-receptor negative breast cancer — then you could use their respective EFO identifiers (EFO_0000305, EFO_1000649, or EFO_1000650) with the function `get_scores()`:

```
scores_bc <- get_scores(efo_id = unique(mavaddat2018_scores@traits[['efo_id']]))
quincunx::n(scores_bc)
#> [1] 96
```

So there are 96 scores present in the PGS Catalog. Included in this set are the 6 scores originating from Mavaddat et al. (2018) (https://doi.org/10.1016/j.ajhg.2018.11.002). If we want to proceed with analysing these other scores without including those from the Mavaddat’s study, we could use the function `setdiff()` to remove them from the S4 `scores` object:

```
# Use quincunx::setdiff to avoid collision with dplyr::setdiff()
bc_scores_not_mavaddat2018 <- quincunx::setdiff(scores_bc, mavaddat2018_scores)
quincunx::n(bc_scores_not_mavaddat2018)
#> [1] 90
bc_scores_not_mavaddat2018@scores[c('pgs_id', 'reported_trait','n_variants')]
#> # A tibble: 90 x 3
#>    pgs_id    reported_trait n_variants
#>    <chr>     <chr>               <int>
#>  1 PGS000001 Breast Cancer          77
#>  2 PGS000028 Breast cancer          83
#>  3 PGS000029 Breast cancer          76
#>  4 PGS000045 Breast cancer          88
#>  5 PGS000050 Breast cancer          44
#>  6 PGS000052 Breast cancer         161
#>  7 PGS000072 Breast cancer         187
#>  8 PGS000153 Breast cancer          66
#>  9 PGS000015 Breast cancer        5218
#> 10 PGS000317 Breast cancer         180
#> # … with 80 more rows
```

For other set operations, check their documentation page: `union()` , `intersect()` and `setequal()` .

### Performance Metrics

According to the PGS Catalog team (https://doi.org/10.1101/2020.05.20.20108217 (https://doi.org/10.1101/2020.05.20.20108217)):

> "Performance Metrics assess the validity of a PGS in a Sample Set, independent of the samples used for score development. Common metrics include standardised effect sizes (odds/hazard ratios [OR/HR], and regression coefficients [Beta]), classification accuracy metrics (e.g. AUROC, C-index, AUPRC), but other relevant metrics (e.g. calibration [Chi-square]) can also be recorded. The covariates used in the model (most commonly age, sex, and genetic principal components (PCs) to account of population structure) are also linking to each set of metrics. Multiple PGS can be evaluated on the same Sample Set and further indexed as directly comparable Performance Metrics."

In this section we will learn how to retrieve details about the evaluation of the polygenic scores developed in Mavaddat *et al.* (2018). To do this, we start by querying the performance metrics for our scores of interest.

Please recall that the `pgs_ids` and `reported_traits` for the polygenic scores newly developed in Mavaddat's publication are:

```
mavaddat2018_scores@scores[c('pgs_id', 'reported_trait')]
#> # A tibble: 6 x 2
#>   pgs_id    reported_trait
#>   <chr>     <chr>
#> 1 PGS000004 Breast Cancer
#> 2 PGS000005 ER-positive Breast Cancer
#> 3 PGS000006 ER-negative Breast Cancer
#> 4 PGS000007 Breast Cancer
#> 5 PGS000008 ER-positive Breast Cancer
#> 6 PGS000009 ER-negative Breast Cancer
```

So we use these PGS identifiers to query the PGS Catalog for performance metrics using the function `get_performance_metrics()`.

```
mavaddat2018_ppm <- get_performance_metrics(pgs_id = mavaddat2018_scores@scores$pgs_i
d)
```

The output is an S4 object with 9 tables:

- performance_metrics, publications, sample_sets, samples, demographics, cohorts, pgs_effect_sizes, pgs_classification_metrics, and pgs_other_metrics.

Reminder | You can quickly check all variables from each table by consulting quincunx cheatsheet (https://github.com/ramiromagno/cheatsheets/blob/master/quincunx/quincunx_cheatsheet.pdf).
Also, a description of each variable is annotated in the `performance_metrics` object help page that can be accessed with `class?performance_metrics`.

#### performance_metrics object | performance_metrics table

In the first table `performance_metrics` we get one performance metrics entity per row, with the following columns: ppm_id, pgs_id, reported_trait, covariates, comments.
Note that all performance metrics are indexed with an unique identifier that starts with "PPM".

```
mavaddat2018_ppm@performance_metrics[1:4]
#> # A tibble: 74 x 4
#>    ppm_id  pgs_id  reported_trait                   covariates
#>    <chr>   <chr>   <chr>                            <chr>
#>  1 PPM000… PGS000… Invasive breast cancer           study, genetic PCs 1-15
#>  2 PPM000… PGS000… Incident breast cancer cases     study, genetic PCs 1-15
#>  3 PPM000… PGS000… Breast cancer intrinsic-like … <NA>
#>  4 PPM000… PGS000… Breast cancer intrinsic-like … <NA>
#>  5 PPM000… PGS000… Breast cancer intrinsic-like … <NA>
#>  6 PPM000… PGS000… Breast cancer intrinsic-like … <NA>
#>  7 PPM000… PGS000… Breast cancer intrinsic-like … <NA>
#>  8 PPM000… PGS000… Metachronous contralateral br… Country
#>  9 PPM000… PGS000… Metachronous contralateral br… Country, age at first diagnos…
#> 10 PPM000… PGS000… Invasive metachronous contral… Country
#> # … with 64 more rows
```

According to the PGS Catalog documentation (http://www.pgscatalog.org/docs/ (http://www.pgscatalog.org/docs/)):

- The `reported_trait` displays the reported trait, often corresponding to the test set names reported in the publication, or more specific aspects of the phenotype being tested (e.g. if the disease cases are incident vs. recurrent events).

- The `covariates` column lists the covariates used in the prediction model to evaluate the PGS. Examples include: age, sex, smoking habits, etc.
- The `comments` column is a field where additional relevant information can be added to aid with understanding a particular performance metrics.

Looking at the `performance_metrics` table, we can see that one polygenic score ( `pgs_id` ) can be associated with several performance metrics ( `ppm_id` ), e.g., PGS000007 associates with 4 PPMs:

```
dplyr::filter(mavaddat2018_ppm@performance_metrics, pgs_id == 'PGS000007')
#> # A tibble: 4 x 5
#>   ppm_id    pgs_id    reported_trait                covariates          comments
#>   <chr>     <chr>     <chr>                         <chr>               <chr>
#> 1 PPM000008 PGS000007 Invasive breast cancer        study, genetic PCs … <NA>
#> 2 PPM000384 PGS000007 Breast Cancer (personal his… age at menarche      <NA>
#> 3 PPM000386 PGS000007 Breast Cancer (personal his… age, sex             <NA>
#> 4 PPM000388 PGS000007 Breast Cancer (personal his… <NA>                 <NA>
```

This means that the polygenic score (PGS000007) has been validated several times, using different models. In this case, we can immediately see that PGS000007 performance has been evaluated, for example, for alternative breast cancer types (different `reported_traits` ), namely:

- Invasive breast cancer, Breast Cancer (personal history).

Additionally, we can see that the same `reported_trait` (*Breast Cancer (personal history)*) has been validated by 3 different performance metrics (PPMs): PPM000384, PPM000386, PPM000388; each having included different covariates in the model: age at menarche, age, sex, NA. (NA means that data for this field is *Not Available* in the records).

#### performance_metrics object | publications table

The publications table is dedicated to hold information related to the publications associated with the performance metrics. The column `pgp_id` links each performance metrics to the respective publication where that performance metrics was reported and collected.

```
mavaddat2018_ppm@publications
#> # A tibble: 74 x 8
#>    ppm_id pgp_id pubmed_id publication_date publication title author_fullname
#>    <chr>  <chr>  <chr>     <date>           <chr>       <chr> <chr>
#>  1 PPM00… PGP00… 30554720  2018-12-13       Am J Hum G… Poly… Mavaddat N
#>  2 PPM00… PGP00… 30554720  2018-12-13       Am J Hum G… Poly… Mavaddat N
#>  3 PPM00… PGP00… 32424353  2020-05-18       Nat Genet   Geno… Zhang H
#>  4 PPM00… PGP00… 32424353  2020-05-18       Nat Genet   Geno… Zhang H
#>  5 PPM00… PGP00… 32424353  2020-05-18       Nat Genet   Geno… Zhang H
#>  6 PPM00… PGP00… 32424353  2020-05-18       Nat Genet   Geno… Zhang H
#>  7 PPM00… PGP00… 32424353  2020-05-18       Nat Genet   Geno… Zhang H
#>  8 PPM00… PGP00… 33022221  2020-10-05       Am J Hum G… Brea… Kramer I
#>  9 PPM00… PGP00… 33022221  2020-10-05       Am J Hum G… Brea… Kramer I
#> 10 PPM00… PGP00… 33022221  2020-10-05       Am J Hum G… Brea… Kramer I
#> # … with 64 more rows, and 1 more variable: doi <chr>
```

Here we can immediately see that there are more publications than just the Mavaddat *et al.* (pubmed_id = 30554720) that we started with.
This is expected because we requested all the performance metrics for the PGS scores that were newly **developed** by Mavaddat *et al.*; but these scores have been subsequently **evaluated** by other posterior publications, and accordingly have performance metrics reported in these posterior evaluations.
This is easily confirmed by checking that all other publications are dated after December 13th 2018, which is the date of publication of the original Mavaddat *et al.* paper.

```
mavaddat2018_ppm@publications$publication_date %>% unique()
#> [1] "2018-12-13" "2020-05-18" "2020-10-05" "2020-07-15" "2020-12-14"
#> [6] "2020-03-12" "2019-11-26"
```

We can choose to look at those publications later to see what evaluations (performance metrics) were reported in them (for which traits, adjusting for which covariates, etc.).

But for now, we are only interested in studying the performance metrics reported by Mavaddat *et al.* for the PGS scores newly developed in this publication. So, lets proceed with creating a vector containing only the `ppm_ids` of interest. We will do this by filtering the `ppm_ids` for the `pubmed_id` corresponding to the Mavaddat publication. We will then use this vector to subset the following tables (to display only the metrics for these PPMs of interest).

```
mavaddat2018_ppm_ids <- mavaddat2018_ppm@publications %>% dplyr::filter(pubmed_id ==
"30554720") %>% dplyr::pull(ppm_id)
mavaddat2018_ppm_ids
#> [1] "PPM000004" "PPM000005" "PPM000006" "PPM000007" "PPM000008" "PPM000009"
#> [7] "PPM000010"

# Find the corresponding pgs_id, reported_trait, covariates, and comments
mavaddat2018_ppm@performance_metrics %>% dplyr::filter(ppm_id %in% mavaddat2018_ppm_i
ds)
#> # A tibble: 7 x 5
#>   ppm_id   pgs_id   reported_trait          covariates        comments
#>   <chr>    <chr>    <chr>                   <chr>             <chr>
#> 1 PPM0000… PGS0000… Invasive breast cancer study, genetic … <NA>
#> 2 PPM0000… PGS0000… Incident breast cance… study, genetic … Included only 306 o…
#> 3 PPM0000… PGS0000… ER-positive breast ca… study, genetic … <NA>
#> 4 PPM0000… PGS0000… ER-negative breast ca… study, genetic … <NA>
#> 5 PPM0000… PGS0000… Invasive breast cancer study, genetic … <NA>
#> 6 PPM0000… PGS0000… ER-positive breast ca… study, genetic … <NA>
#> 7 PPM0000… PGS0000… ER-negative breast ca… study, genetic … <NA>
```

Here, we can immediately see that the Mavaddat publication has evaluated the PGS000004 twice, with two performance metrics (PPM000004 and PPM000005), that are different because they include a different set of SNPs (see the `comments` column for PPM000005).

#### performance_metrics object | sample_sets table

The PGS Catalog provides a Sample Set Id (PSS) that links the PPMs to the sample sets that were used to evaluate the corresponding PGS. This mapping is stored in the `sample_sets` table.

```
mavaddat2018_ppm@sample_sets %>% dplyr::filter(ppm_id %in% mavaddat2018_ppm_ids)
#> # A tibble: 7 x 2
#>   ppm_id    pss_id
#>   <chr>     <chr>
#> 1 PPM000004 PSS000004
#> 2 PPM000005 PSS000007
#> 3 PPM000006 PSS000005
#> 4 PPM000007 PSS000006
#> 5 PPM000008 PSS000004
#> 6 PPM000009 PSS000005
#> 7 PPM000010 PSS000006
```

#### performance_metrics object | samples table

The `samples` table gathers more relevant information regarding the samples used for the relevant evaluations. This table contains the following columns:

- ppm_id, pss_id, sample_id, stage, sample_size, sample_cases, sample_controls, sample_percent_male, phenotype_description, ancestry, ancestry_description, ancestry_country, ancestry_additional_description, study_id, pubmed_id, cohorts_additional_description.

Please note that the samples are not identified in PGS Catalog with a global unique identifier, but *quincunx* assigns a surrogate identifier ( `sample_id` ) to allow the mapping between tables.

```
mavaddat2018_ppm@samples %>% dplyr::filter(ppm_id %in% mavaddat2018_ppm_ids)
#> # A tibble: 7 x 16
#>   ppm_id pss_id sample_id stage sample_size sample_cases sample_controls
#>   <chr>  <chr>      <int> <chr>       <int>        <int>           <int>
#> 1 PPM00… PSS00…         1 eval…       29751        11428           18323
#> 2 PPM00… PSS00…         1 eval…      190040         3215          186825
#> 3 PPM00… PSS00…         1 eval…       11428         7992            3436
#> 4 PPM00… PSS00…         1 eval…       11428         1259           10169
#> 5 PPM00… PSS00…         1 eval…       29751        11428           18323
#> 6 PPM00… PSS00…         1 eval…       11428         7992            3436
#> 7 PPM00… PSS00…         1 eval…       11428         1259           10169
#> # … with 9 more variables: sample_percent_male <dbl>,
#> #   phenotype_description <chr>, ancestry <chr>, ancestry_description <chr>,
#> #   ancestry_country <chr>, ancestry_additional_description <chr>,
#> #   study_id <chr>, pubmed_id <chr>, cohorts_additional_description <chr>
```

Here we can, for example see that the sample sizes are very different for each evaluation. Let's take a quick look at their values.

```
mavaddat2018_ppm@samples %>%
  dplyr::filter(ppm_id %in% mavaddat2018_ppm_ids) %>%
  dplyr::select(ppm_id, sample_cases, sample_controls) %>%
  tidyr::pivot_longer(!ppm_id, names_to = "sample_type", values_to = "count") %>%
  ggplot2::ggplot(ggplot2::aes(fill=sample_type, y=count, x=ppm_id)) +
  ggplot2::geom_bar(position="dodge", stat="identity") +
  ggplot2::theme(axis.text.x = ggplot2::element_text(angle = 20, vjust = 0.5, hjust=
0.5))
```

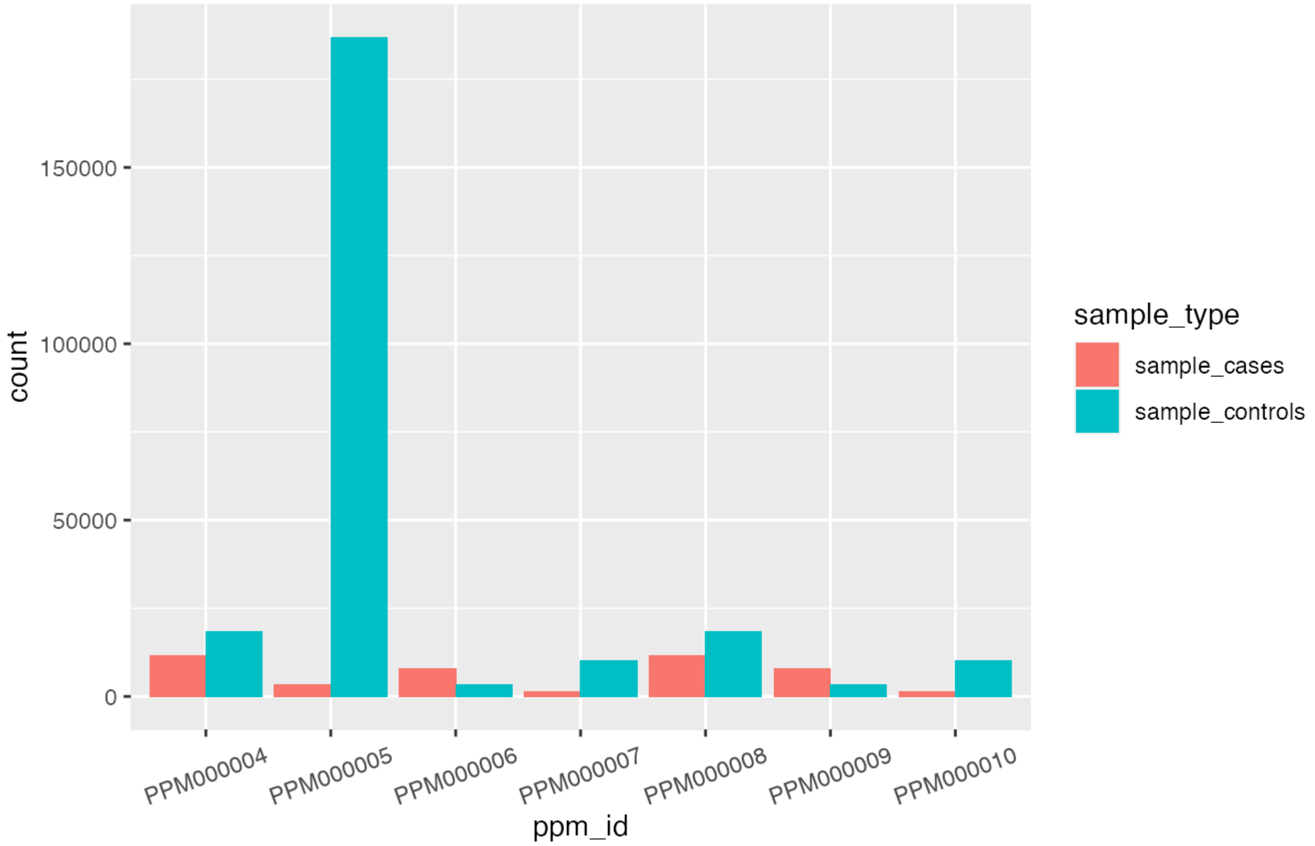


This plot clearly shows that PPM000005 uses a very large sample (particularly regarding the number of controls) when compared with all other reported PPMs.

#### performance_metrics object | demographics table

The `demographics` table holds information regarding the demographics' variables of each `sample`. Each demographics' variable (row) is uniquely identified by the combination of values from the columns: `ppm_id`, `pss_id`, `sample_id`, and `variable`. Currently, the PGS Catalog only describes two demographic variables: *age of participants* and *follow-up time*.

The columns presented in the table are:

- ppm_id, pss_id, sample_id, variable, estimate_type, estimate, unit, variability_type, variability, interval_type, interval_lower, interval_upper

```
mavaddat2018_ppm@demographics %>% dplyr::glimpse()
#> Rows: 2
#> Columns: 12
#> $ ppm_id           <chr> "PPM001345", "PPM001347"
#> $ pss_id           <chr> "PSS000450", "PSS000450"
#> $ sample_id        <int> 1, 1
#> $ variable         <chr> "age", "age"
#> $ estimate_type    <chr> "mean age (at the end of follow-up)", "mean age (at …
#> $ estimate         <dbl> 58.5, 58.5
#> $ unit             <chr> "years", "years"
#> $ variability_type <chr> NA, NA
#> $ variability      <dbl> NA, NA
#> $ interval_type    <chr> "iqr", "iqr"
#> $ interval_lower   <dbl> 45.1, 45.1
#> $ interval_upper   <dbl> 72.2, 72.2
```

Here we can see that neither of the PPMs shown is from the Mavaddat paper. This means that the performance metrics reported in the paper do not have any information regarding the demographics' variables that belong in this table (age and follow-up time).

#### performance_metrics object | Cohorts table

Similarly to the `cohorts` table described above (in the `scores` object class), this table shows which cohorts are associated with each sample. However here, the unique identification of a sample can be obtained by combining the values of the first three columns: `ppm_id` , `pss_id` and `sample_id` .

```
mavaddat2018_ppm@cohorts %>% dplyr::filter(ppm_id %in% mavaddat2018_ppm_ids)
#> # A tibble: 61 x 5
#>    ppm_id   pss_id  sample_id cohort_symbol cohort_name
#>    <chr>    <chr>       <int> <chr>         <chr>
#>  1 PPM0000… PSS000…         1 AHS           Agricultural Health Study
#>  2 PPM0000… PSS000…         1 BGS           Breakthrough Generations Study
#>  3 PPM0000… PSS000…         1 EPIC          European Prospective Investigation …
#>  4 PPM0000… PSS000…         1 FHRISK        Family History Clinic in Manchester
#>  5 PPM0000… PSS000…         1 KARMA         UNKNOWN
#>  6 PPM0000… PSS000…         1 NHS           Nurses Health Study
#>  7 PPM0000… PSS000…         1 NHS2          Nurses Health Studies II
#>  8 PPM0000… PSS000…         1 PLCO          Prostate, Lung, Colorectal, and Ova…
#>  9 PPM0000… PSS000…         1 PROCAS        Breast Screening Programme (NHSBSP)…
#> 10 PPM0000… PSS000…         1 SISTER        UNKNOWN
#> # … with 51 more rows
```

#### performance_metrics object | PGS effect sizes & PGS classification metrics & PGS other metrics tables

The three final tables of the `performance_metrics` object hold the performance metrics themselves used in the validation. Each table presents the same column structure with 11 total columns, where the second column is different between the three tables. This column shows an id created by *quincunx* that is used to identify each of the individual metrics tables ( `effect_size_id` , `classification_metrics_id` , or `other_metrics_id` ). All other columns are for the same variables:

- ppm_id, effect_size_id, estimate_type_long, estimate_type, estimate, unit, variability_type, variability, interval_type, interval_lower, interval_upper.

(These column names are specifically from the `pgs_effect_sizes` table, as indicated by the second column name. All other columns are equally named in all three tables).

According to the PGS Catalog online documentation (http://www.pgscatalog.org/docs/ (http://www.pgscatalog.org/docs/)):

> “The reported values of the performance metrics are all reported similarly (e.g. the estimate is recorded along with the 95% confidence interval (if supplied)) and grouped according to the type of statistic they represent:
> - **PGS Effect Sizes (per SD change)** | Standardized effect sizes, per standard deviation [SD] change in PGS. Examples include regression coefficients (Betas) for continuous traits, Odds ratios (OR) and/or Hazard ratios (HR) for dichotomous traits depending on the availability of time-to-event data.
> - **PGS Classification Metrics** | Examples include the Area under the Receiver Operating Characteristic (AUROC) or Harrell’s C-index (Concordance statistic).
> - **Other Metrics** | Metrics that do not fit into the other two categories. Examples include: R2 (proportion of the variance explained), or reclassification metrics.”

Now, lets briefly explore the data in these tables (as usual filtered for only the PPMs that were newly reported in Mavaddat *et al.*).

```
mavaddat2018_ppm@pgs_effect_sizes %>% dplyr::filter(ppm_id %in% mavaddat2018_ppm_ids)
#> # A tibble: 7 x 11
#>   ppm_id effect_size_id estimate_type_l… estimate_type estimate unit
#>   <chr>           <int> <chr>            <chr>            <dbl> <chr>
#> 1 PPM00…              1 Odds Ratio       OR                1.61 <NA>
#> 2 PPM00…              1 Hazard Ratio     HR                1.59 <NA>
#> 3 PPM00…              1 Odds Ratio       OR                1.68 <NA>
#> 4 PPM00…              1 Odds Ratio       OR                1.45 <NA>
#> 5 PPM00…              1 Odds Ratio       OR                1.66 <NA>
#> 6 PPM00…              1 Odds Ratio       OR                1.73 <NA>
#> 7 PPM00…              1 Odds Ratio       OR                1.44 <NA>
#> # … with 5 more variables: variability_type <chr>, variability <dbl>,
#> #   interval_type <chr>, interval_lower <dbl>, interval_upper <dbl>
mavaddat2018_ppm@pgs_classification_metrics %>% dplyr::filter(ppm_id %in% mavaddat201
8_ppm_ids)
#> # A tibble: 6 x 11
#>   ppm_id classification_… estimate_type_l… estimate_type estimate unit
#>   <chr>             <int> <chr>            <chr>            <dbl> <chr>
#> 1 PPM00…                1 Area Under the … AUROC            0.63  <NA>
#> 2 PPM00…                1 Area Under the … AUROC            0.641 <NA>
#> 3 PPM00…                1 Area Under the … AUROC            0.601 <NA>
#> 4 PPM00…                1 Area Under the … AUROC            0.636 <NA>
#> 5 PPM00…                1 Area Under the … AUROC            0.647 <NA>
#> 6 PPM00…                1 Area Under the … AUROC            0.6   <NA>
#> # … with 5 more variables: variability_type <chr>, variability <dbl>,
#> #   interval_type <chr>, interval_lower <dbl>, interval_upper <dbl>
mavaddat2018_ppm@pgs_other_metrics %>% dplyr::filter(ppm_id %in% mavaddat2018_ppm_ids
)
#> # A tibble: 0 x 11
#> # … with 11 variables: ppm_id <chr>, other_metrics_id <int>,
#> #   estimate_type_long <chr>, estimate_type <chr>, estimate <dbl>, unit <chr>,
#> #   variability_type <chr>, variability <dbl>, interval_type <chr>,
#> #   interval_lower <dbl>, interval_upper <dbl>
```

We can immediately see that the third table (reserved for metrics other than effect sizes and classification metrics) is empty; and that the effect sizes were estimated using Odds Ratio and Hazard Ratio, and the classification metric applied was AUROC.

Now we can look at the reported values for each PPM metrics and decide if the validation is relevant, and therefore make an informed choice to use the associated PGS score for our own study, and eventually apply it to our own dataset.

### Concluding remarks

Useful reminders:

- You can quickly check all variables from each table by consulting quincunx cheatsheet (https://github.com/ramiromagno/cheatsheets/blob/master/quincunx/quincunx_cheatsheet.pdf).
- All *quincunx* functions are annotated with help pages accessed with `?function_name` (e.g. `?get_scores`, `?get_traits`, `?get_performance_metrics`).
- All tables retrieved by *quincunx's* functions are annotated with help files that describe all variables present. These are accessible via `class?object_name` (e.g. `class?scores`, `class?performance_metrics`).
- The *quincunx* package has been published in a peer-reviewed journal. Use `citation(package="quincunx")` to get the full paper citation.

---
