## Additional File 2 - quincunx cheatsheet for "quincunx: an R package to query, download and wrangle PGS Catalog data"

### PGS Catalog access with quincunx

#### Introduction

The **PGS Catalog** is a service provided by the EMBL-EBI and University of Cambridge that offers a manually curated and freely available database of published polygenic scores (PGS): `https://www.pgscatalog.org/`.

The PGS Catalog data provided by the **REST API** is organised around five core entities:

- PGS **Polygenic Scores**
- PGP **PGS Publications**
- PSS **PGS Sample Sets**
- PPM **PGS Performance Metrics**
- EFO **EFO traits**

#### Get PGS Catalog Entities

**quincunx** facilitates the access to the Catalog via the REST API, allowing you to programmatically retrieve data directly into R. Each of the five entities is mapped to an S4 object of a class of the same name.

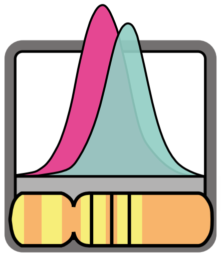

| PGS CATALOG | RETRIEVAL FUNCTIONS | S4 CLASSES |
|---|---|---|
| | `get_scores()` | scores |
| | `get_publications()` | publications |
| | `get_sample_sets()` | sample_sets |
| | `get_performance_metrics()` | performance_metrics |
| | `get_traits()` | traits |

Query criteria for retrieval functions, e.g., PGS can be queried by either `pgs_id`, `efo_id` or `pubmed_id`. These correspond to the criteria exposed by the PGS Catalog REST API: `https://www.pgscatalog.org/rest/`.

| Search by | Example | PGS | PGP | PSS | PPM | EFO |
|---|---|---|---|---|---|---|
| pgs_id | "PGS000001" | ■ | ■ | ■ | ■ | |
| pgp_id | "PGP000001" | | ■ | | | |
| pss_id | "PSS000001" | | | ■ | | |
| ppm_id | "PPM000001" | | | | ■ | |
| efo_id | "EFO_0000249" | ■ | | | | ■ |
| pubmed_id | "25855707" | ■ | ■ | | | |
| author | "Mavaddat" | | ■ | | | |
| trait_term | "Alzheimer" | | | | | ■ |

#### PGS Catalog Entities in R

PGS Catalog entities are represented as S4 classes in R. Each class represents a relational database of tidy data tables. All objects start with a table with the same name as the class. Combination of variables indicated in bold renders each row unique in each table.

##### S4 class scores

`scores`
- **pgs_id**
- pgs_name
- scoring_file
- matches_publication
- reported_trait
- trait_additional_description
- pgs_method_name
- pgs_method_params
- n_variants
- n_variants_interactions
- assembly
- license
- beta_unit

`samples`
- **pgs_id**
- **sample_id**
- stage
- sample_size
- sample_cases
- sample_controls
- sample_percent_male
- phenotype_description
- ancestry
- ancestry_description
- ancestry_country
- ancestry_additional_description
- study_id
- pubmed_id
- cohorts_additional_description

`demographics`
- **pgs_id**
- **sample_id**
- **variable**
- estimate_type
- estimate
- unit
- variability_type
- variability
- interval_type
- interval_lower
- interval_upper

`publications`
- **pgs_id**
- **pgp_id**
- pubmed_id
- publication_date
- publication
- title
- author_fullname
- doi

`traits`
- **pgs_id**
- **efo_id**
- trait
- description
- url

`cohorts`
- **pgs_id**
- **sample_id**
- **cohort_symbol**
- cohort_name

##### S4 class publications

`publications`
- **pgp_id**
- pubmed_id
- publication_date
- publication
- title
- author_fullname
- doi
- authors

`pgs_ids`
- **pgp_id**
- **pgs_id**
- stage

##### S4 class traits

`traits`
- **efo_id**
- **parent_efo_id**
- is_child
- trait
- description
- url

`pgs_ids`
- **efo_id**
- **parent_efo_id**
- is_child
- pgs_id

`child_pgs_ids`
- **efo_id**
- **parent_efo_id**
- is_child
- child_pgs_id

3x `trait_{categories, synonyms, mapped_terms}`
- **efo_id**
- **parent_efo_id**
- is_child
- trait_{category, synonyms, mapped_terms}

##### S4 class sample_sets

`sample_sets`
- **pss_id**
- pgs_name
- scoring_file
- matches_publication
- reported_trait
- trait_additional_description
- pgs_method_name
- pgs_method_params
- n_variants
- n_variants_interactions
- assembly
- license
- beta_unit

`cohorts`
- **pss_id**
- **sample_id**
- **cohort_symbol**
- cohort_name

`samples`
- **pss_id**
- **sample_id**
- stage
- sample_size
- sample_cases
- sample_controls
- sample_percent_male
- phenotype_description
- ancestry
- ancestry_description
- ancestry_country
- ancestry_additional_description
- study_id
- pubmed_id
- cohorts_additional_description

`demographics`
- **pss_id**
- **sample_id**
- **variable**
- estimate_type
- estimate
- unit
- variability_type
- variability
- interval_type
- interval_lower
- interval_upper

##### S4 class performance_metrics

`performance_metrics`
- **ppm_id**
- pgs_id
- reported_trait
- covariates
- comments

`publications`
- **ppm_id**
- **pgp_id**
- pubmed_id
- publication_date
- publication
- title
- author_fullname
- doi

`sample_sets`
- **ppm_id**
- **pss_id**

`samples`
- **ppm_id**
- **pss_id**
- **sample_id**
- stage
- sample_size
- sample_cases
- sample_controls
- sample_percent_male
- phenotype_description
- ancestry
- ancestry_description
- ancestry_country
- ancestry_additional_description
- study_id
- pubmed_id
- cohorts_additional_description

`demographics`
- **ppm_id**
- **pss_id**
- **sample_id**
- **variable**
- estimate_type
- estimate
- unit
- variability_type
- variability
- interval_type
- interval_lower
- interval_upper

`cohorts`
- **ppm_id**
- **pss_id**
- **sample_id**
- **cohort_symbol**
- cohort_name

3x `pgs_{effect_sizes,classification_metrics,other_metrics}`
- **ppm_id**
- **{effect_size_id, classification_metrics_id, other_metrics_id}**
- estimate_type_long
- estimate_type
- estimate
- unit
- variability_type
- variability
- interval_type
- interval_lower
- interval_upper

CC BY SA Ramiro Magno • Learn more at https://maialab.org/quincunx • quincunx version 0.0.1 • Updated: 2021-02-19

### Other S4 Entities

Besides the five PGS Catalog entities, there are three other objects that can be retrieved from the REST API: `trait_categories`, `cohorts` and `releases`.

##### S4 class trait_categories

`trait_categories`
- **trait_category**

`trait`
- **trait_category**
- **efo_id**
- trait
- description
- url

##### S4 class cohorts

`cohorts`
- **cohort_symbol**
- cohort_name

`pgs_ids`
- **cohort_symbol**
- **pgs_id**
- stage

##### S4 class releases

`releases`
- **date**
- n_pgs
- n_ppm
- n_pgp
- notes

3x `{pgs_ids, ppm_ids,pgp_ids}`
- **date**
- **{pgs_id, ppm_id, pgp_id}**

### PGS Construction Process

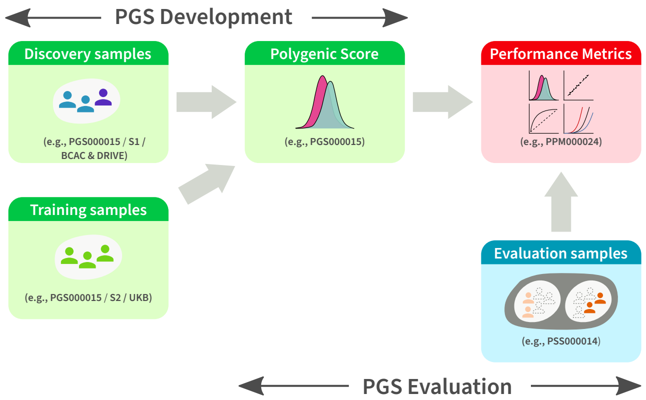


Samples and Polygenic Scores (PGS) are annotated according to their utilisation context in the PGS construction process, i.e. the `stage` variable in quincunx:

- `"discovery"`
- `"training"`
- `"development"` (`"discovery"` or `"training"`)
- `"evaluation"`

### Cohorts, Samples and Sample Sets

##### Cohorts

A cohort is a group of individuals with a shared characteristic. Cohorts are identified in quincunx by the `cohort_symbol` variable.

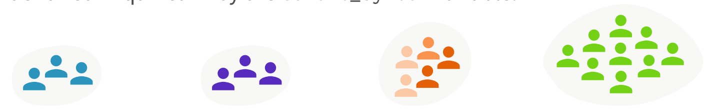


##### Samples

A sample is a group of participants associated with none, one or more catalogued cohorts. The selection from a cohort can be either a subset or its totality. Samples are not identified in PGS Catalog with a global unique identifier, but quincunx assigns a surrogate identifier (`sample_id`) to allow relations between tables.

Possible compositions of samples:

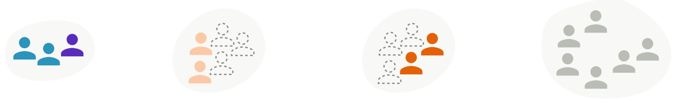


##### Sample Sets

A sample set is a group of samples **used in a polygenic score evaluation.** Each sample set is identified in the PGS Catalog by a unique sample set identifier (PSS ID).

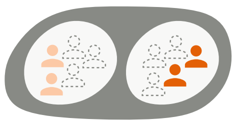

A sample set of two samples.
(e.g., PSS000020: S1 and S2)

### Manipulate Cases of S4 Entities

Get a **scores** object **s** consisting of two polygenic scores (PGS):

```
s <- get_scores(pgs_id = c('a', 'b'))
```

Subset object **s** by either identifier or position using `` `[` ``:

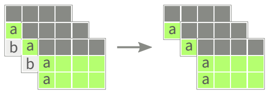


```
s['a'] # Subset by identifier
```

```
s[1] # Subset by position
```

Combine two scores' objects:

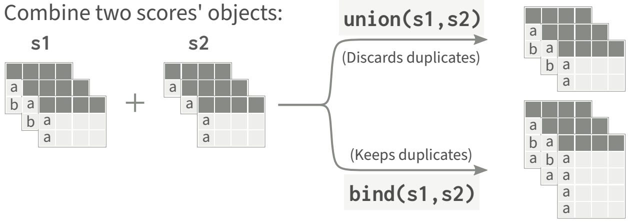


### Polygenic scoring file

PGS scoring files are provided by the PGS Catalog to allow computation of polygenic scores by users. These files are hosted at the PGS Catalog FTP server: `http://ftp.ebi.ac.uk/pub/databases/spot/pgs/scores/`. They are labelled by their respective PGS Score ID (e.g. `PGS000001.txt.gz`). For more details please visit: `https://maialab.org/quincunx/articles/pgs-scoring-file.html`.

##### File Format

Each scoring file contains variant identification, effect alleles and respective score weights. The file is formatted as a gzipped tab-delimited text file, with a header containing brief metada about the score. You can read PGS scoring files into R with `read_scoring_file()`.

`PGS000117.txt.gz`

```
### PGS CATALOG SCORING FILE - see www.pgscatalog.org/downloads/#dl_ftp for…
## POLYGENIC SCORE (PGS) INFORMATION
# PGS ID = PGS000117
# Reported Trait = Cardiovascular Disease
# Original Genome Build = GRCh37
# Number of Variants = 267863
## SOURCE INFORMATION
# PGP ID = PGP000054
# Citation = Elliott J et al. JAMA (2020). doi:10.1001/jama.2019.22241
rsID       chr_name chr_position effect_allele reference_allele effect_weight
rs11240779 1        808631       A             G                0.00077622
rs1921     1        949608       A             G                -0.00583829
rs2710890  1        958905       G             A                -0.00182583
rs4970349  1        967658       T             C                -0.001855691
…
```

##### Columns

The following table lists all possible columns in a PGS scoring file. A few columns are required (R), and most are optional (O); either the `rsID` alone or the combination of `chr_name` and `chr_position` are required, with the other being optional.

| Column (Requirement) | Description | Example |
|---|---|---|
| rsID (R/O) | dbSNP Accession ID | "rs554219" |
| chr_name (R/O) | Chromosome name | "11" |
| chr_position (R/O) | Chromosome position | 69516874 |
| effect_allele (R) | Effect allele | "G" |
| reference_allele (O) | Reference allele | "C" |
| effect_weight (R) | Variant weight | 0.117 |
| locus_name (O) | Locus name | "CCND1" |
| weight_type (O) | Type of weight | "log(OR)", "beta_cox" |
| allelefrequency_effect (O) | Effect allele frequency | 0.410 |
| is_interaction (O) | Variant interaction? | TRUE or FALSE |
| is_recessive (O) | Recessive inheritance model? | TRUE or FALSE |
| is_haplotype (O) | Is effect allele a haplotype? | TRUE or FALSE |
| is_diplotype (O) | Is effect allele a diplotype? | TRUE or FALSE |
| imputation_method (O) | Imputation method | TODO |
| variant_description (O) | Variant description | TODO |
| inclusion_criteria (O) | Score inclusion criteria | TODO |
| OR (O) | Odds Ratio | 1.12 |
| HR (O) | Hazard Ratio | 1.08 |

CC BY SA Ramiro Magno • Learn more at https://maialab.org/quincunx • quincunx version 0.0.1 • Updated: 2021-02-19
